# Exploring use of ozone nanobubbles for removal of cyanobacteria and co-occurring antimicrobial resistance genes in water supply and reuse systems

**DOI:** 10.1101/2025.10.15.682302

**Authors:** Ushalini Saththiyananthan, Calum J. Walsh, Steven Newham, Mark Putman, Daniel Flanagan, Karen Rouse, Filippo Nelli, Elnaz Karamati Niaragh, Louise Judd, Karolina Mercoulia, Torsten Seemann, Ming Su, Min Yang, Linda Blackall, Benjamin P Howden, Eric Wert, Arash Zamyadi

## Abstract

Harmful cyanobacterial blooms present persistent risks to both drinking water security and wastewater reuse, driving the need for advanced treatment strategies. Treatment barrier(s) need to be capable of simultaneously controlling cyanobacteria, cyanotoxins, and co-occurring contaminants like bloom-associated antimicrobial resistance genes (ARGs) particularly in the case of recycling treated wastewater. Ozone nanobubble technology has emerged as a promising innovation, offering extended oxidative stability and enhanced interfacial reactivity compared to conventional ozonation. Hence this research objectives were to (a) assess the removal performance of ozone nanobubbles in eliminating cyanobacteria and their co-occurring contaminants in comparison to conventional ozone systems, and (b) investigate the repeatability of the results in varying background water qualities, ozone decay and the potential for by-products formation

Ozonation using nanobubbles enhanced oxidation performance by 19%-34% in the drinking water reservoir compared to conventional ozonation while keeping the ozone concentration below 2mg/l. Lower oxidation efficiencies were observed in treated wastewater compared to drinking water sources, reflecting the higher content of organic matter and suspended solids, and oxidant demand characteristic of recycled water systems. Despite these challenges, ozone nanobubbles consistently outperformed conventional ozonation in reducing both cyanobacterial biomass and cell viability, underscoring their potential as an advanced “polishing” step for algal management in wastewater reuse applications.

By exploring fate of ARGs alongside cyanobacteria and toxin removal, this work extends beyond traditional ozonation trials. It provides valuable field-based evidence that bridges the divide between laboratory efficacy and full-scale operational performance. Future studies should build on this by exploring combined or sequential treatment barriers that enhance DNA degradation, thereby addressing both cellular and genetic risks in water supply and reuse systems.

Observing the action of nanobubbles under dynamic, real-world water quality conditions is currently challenging; however, this study’s novel field trials demonstrate potential nanobubble applications and provide valuable insights to guide future investigations. The results reinforce the broader applicability of ozone nanobubble technology for multi-target contaminant control in water reservoirs.

## 1. Introduction

Cyanobacteria are photoautotrophs prevalent in aquatic ecosystems (Chorus and Welker, 2021). However, due to the effects of climate change, increase in human activities caused by population growth, and accelerated eutrophication, the frequency of intensive cyanobacterial growth events, called blooms, is increasing in many natural and constructed water bodies worldwide (Paerl and Paul, 2012; Krienitz et al., 2013; Chorus and Welker, 2021). Cyanobacteria produce a broad spectrum of metabolites, including cyanotoxins, namely microcystins, cylindrospermopsins, anatoxins, saxitoxins, and potentially beta-N-methylamino-L-alanine (BMAA), as well as taste and odour (T&O) compounds such as geosmin and 2-methylisoborneol (2-MIB), that impact water quality and consumer perception (Su et al., 2015; Merel et al., 2013; Chorus and Welker, 2021; Kibuye et al., 2021a). Human health impacts of cyanotoxins include acute liver damage, gastrointestinal disturbances, serious neurotoxicosis, respiratory problems and allergic reactions (Chorus & Welker, 2021). Meanwhile, non-toxic T&O compounds deteriorate the aesthetic quality of drinking water even in concentrations as low as 7 ng/L (Chen et al., 2019; Kibuye et al., 2021a). These harmful and nuisance blooms are affecting drinking water reservoirs and treated wastewater stabilization ponds scheduled for reuse, posing a major risk to water availability (Coggins et al., 2019; Chorus and Welker, 2021; Kibuye et al., 2021a, 2021b and 2021c; Romanis et al., 2023; Romanis et al., 2024).

Cyanobacterial blooms in treated wastewater stabilization ponds leads to their potential extensive contact with antimicrobial resistance genes (ARG), potential human and animal pathogens, and other bacteria associated with treated wastewater (Wang et al., 2022; Zhou 2023). Some studies have linked cyanobacterial blooms within natural and constructed water systems to prevalence of ARGs, particularly blooms of taxa such as *Synechococcus, Microcystis*, and, *Planktothrix* (Zhang et al., 2020; Wang et al., 2020; Wang et al., 2021; Li et al., 2021; Wang et al., 2022; Volk et al., 2023; Ji et al., 2024). While proposed mechanisms linking cyanobacterial booms to ARG proliferation require further investigation to understand the underlying drivers, these blooms must be managed with a dual treatment approach in mind: targeting both cyanobacteria and associated ARGs (Volk et al., 2023; Ji et al., 2024).

A variety of physical, chemical, and biological processes are widely used in both drinking water and treated wastewater reuse systems to remove cyanobacterial cells and their associated contaminants (Chorus and Welker, 2021; Kibuye et al., 2021a, 2021b and 2021c). However, each of these approaches faces limitations, particularly in effectively addressing both intact cells, colonies, and associated chemical and biological contaminants (Zamyadi et al., 2019; Kibuye et al., 2021a, 2021b and 2021c; Su et al., 2021; Zamyadi and Blackall, 2024). A major concern in both drinking water and reuse contexts is the occurrence of breakthrough events. During such event cells and/or their associated contaminants pass through the treatment barriers. These events compromise achievement of water and public health safety guidelines, treatment reliability, and regulatory compliance (Zamyadi et al., 2012; Zamyadi et al., 2019; Zamyadi and Blackall, 2024).

Given these challenges, it is increasingly clear that targeting cyanobacteria and their associated contaminants at early treatment stages or pre-treatment, before they enter the main treatment process, is advisable. Use of oxidising agents, like H_2_O_2_, KMnO_4_, Cl_2_, and O_3_, as a pre-treatment control strategy offers the potential to minimize the cyanobacterial load and associated contaminants early in the process, reducing downstream treatment challenges and potentially preventing breakthrough events altogether (Kibuye et al., 2021a, 2021b and 2021c; Zamyadi and Blackall, 2024). The overall effectiveness of treatment with oxidising agents varies significantly depending on the type of oxidant, the specific contaminants targeted, and operational conditions such as dose, contact time, pH, and water matrix characteristics, and extent to which their use leads to the formation of harmful disinfection by-products (DBPs) (Chorus and Welker, 2021; Kibuye et al., 2021b).

Among oxidants, ozone (O_3_) offers a strong option, providing broad-spectrum reactivity and high potential for the simultaneous degradation of microbial cells, toxins, and emerging contaminants—making it a leading candidate for advanced water and wastewater treatment applications (Coral et al., 2013; Zamyadi et al., 2015; Ikehata and Li, 2018; Kibuye et al., 2021b; Morrison et al., 2021). The ozonation process produces reactive oxygen species (ROS), with hydroxyl radicals (*O H*^•^) being the strongest ROS, capable of effectively degrading the cyanotoxins into non-toxic end products (Manasfi, 2021; Morrison et al., 2021). Conventional ozone treatment at efficient dosages (for example above 5mg/L) is energy intensive and include operational difficulties due to ozone half-life in water and O_3_ losses due to off-gassing (Coral et al., 2013; Zamyadi et al., 2015; Manasfi, 2021; Yang et al., 2023). Conventional systems require high-pressure injection, large gas volumes, and extensive contact infrastructure, making them less feasible for decentralized, energy-conscious and/or regional applications such as wastewater reuse and reservoir management. Also, some cyanobacteria like *Microcystis* tend to form extensive colonies in real-world setting compared to laboratory cultures (Zamyadi and Blackall, 2024); hence it is hypothesized that increasing ozone contact surface area would significantly benefit the removal process by enhancing the breaking of colonies and then oxidizing the individual cells depending on oxidant residual.

Ozone nanobubble technology potentially offers a unique and transformative compared to conventional oxidant-based treatment processes (Kuhn et al., 2025). Nanobubbles are gas filled cavities with diameter of less than 100 nm, which exhibit several unique properties that have been utilized in water and wastewater treatment such as flotation, membrane cleaning, adsorption, aeration, anaerobic digestion and advanced oxidation (Jia et al., 2023; Singh et al., 2024; Soyluoglu et al. 2021, 2022 and 2023; Koundle et al., 2024). The larger surface area of nanobubbles increases gas-liquid mass transfer rate, elevating ozone dissolution in water. By generating ultra-fine, stable bubbles with enhanced gas transfer efficiency and prolonged oxidative activity in water, ozone nanobubble technology is hypothesized to achieve greater contaminant degradation at lower ozone dosages and with significantly reduced energy input (Atkinson et al. 2019; Batagoda et al., 2019; Jia et al., 2023; Singh et al., 2024; Kuhn et al., 2025). This potentially makes them particularly suited for source-critical-point intervention, enabling more sustainable and targeted control of cyanobacterial threats in both drinking water and treated wastewater systems (Soyluoglu et al. 2021, 2022, 2023 and 2025; Chaffin et al., 2024).

There remains a critical need to accurately quantify both the nanobubble population and the trace levels of residual ozone, in ng/L range, to better understand their stability, oxidative potential, and contribution to treatment performance under real-world conditions (Kuhn et al., 2025). Detecting nanobubbles directly in turbid water is technically challenging due to light scattering interference from suspended solids and organic matter, which masks their optical signal (Jia et al., 2023; Singh et al., 2024; Soyluoglu et al. 2021, 2022 and 2023; Koundle et al., 2024). In such conditions, treatment performance indicators, such as microbial inactivation, cyanobacterial cell lysis, and changes in water quality, might serve as practical, indirect evidence of nanobubble presence and effectiveness. This hypothesis requires detailed investigation particularly evidence of repeatability.

This research investigates the application of ozone nanobubble technology as an advanced oxidation barrier for the simultaneous removal and destruction of cyanobacterial blooms and their associated toxins, and, in the context of treated wastewater, for the mitigation of ARGs. The study aims to evaluate ozone nanobubble performance under real-world water supply and reuse conditions, providing mechanistic and operational insights into its potential as a sustainable, multi-contaminant treatment solution. Hence this research objectives were to (a) assess the removal performance of ozone nanobubbles in eliminating cyanobacteria and their co-occurring contaminants in comparison to conventional ozone systems, and (b) investigate the repeatability of the results in varying background water qualities, ozone decay and the potential for by-products formation—a key drawback of ozonation. To the best of authors’ knowledge this research investigates for the first time use of ozone nanobubble for removal of cyanobacterial bloom, cyanotoxins, and ARGs in real-world water supply and reuse systems.

## 2. Material and methods

### 2.1. Water sampling and trial sites

The sampling campaign spanned multiple water types and treatment contexts, enabling comparison of ozone nanobubble performance across contrasting environmental and operational conditions. These sites (Figure 1) include

**Figure 1.**
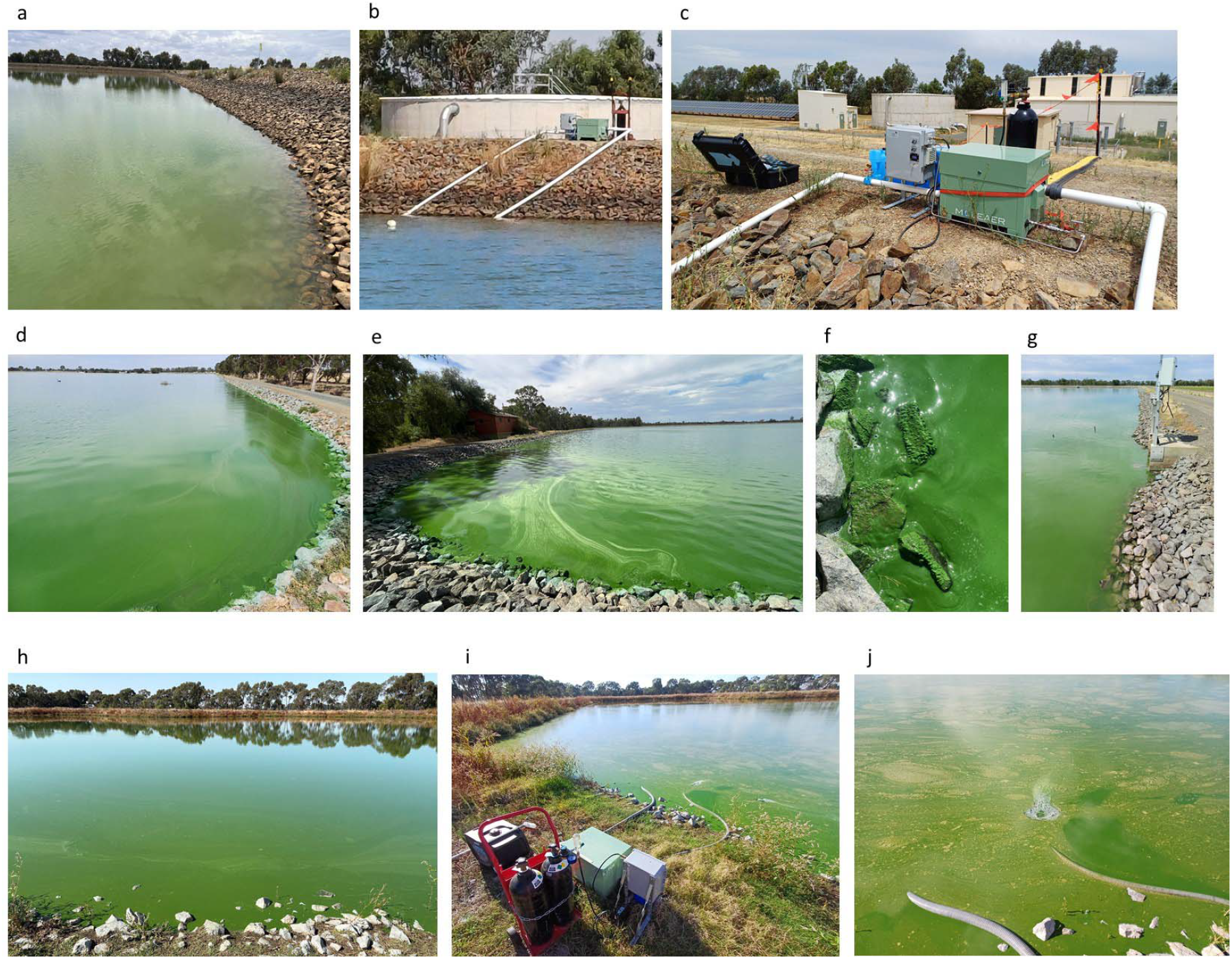
Full-scale experimental sites: (a, b, c) drinking water reservoir; (d, e, f, g) WWTP-post-aeration and Class-C recycled water; and (h, i, j) treated r in lagoon-based system.

i. an off-stream drinking water reservoir studied during seven trial events from beginning to end of the bloom season: 18 November, 6 January, 9 January, 23 January, 4 February, 14 February, and 7 March;
ii. treated wastewater post aeration ponds, labelled “WWTP-aeration-pond”, and treated wastewater at the end of storage ponds before recycled for farming labelled “Class-C”, studied during four trial events: 11 November (T1), 17 December (T2), 6 January (T3), and 4 February (T4), and
iii. a lagoon-based wastewater treatment plant studied during two trial events: 11 April and 16 April.

All these sites are located in southeast of Australia.

The timing of installations and sampling across these sites was coordinated to capture both baseline conditions and treatment effects during bloom-prone months. Figures 1a-1c show the off-stream drinking water reservoir site, where ozone nanobubble and conventional ozone treatments were trialled. Water sampling at this site was undertaken at multiple intervals following dosing, to capture short-term oxidative effects and explore reproducible results across the summer season. Figures 1d–1g represent the wastewater treatment plant sampling points labelled “WWTP-aeration-pond” and “Class-C” recycled water storages, where biomass loads and cyanobacterial densities were substantially higher than in the drinking water reservoir. Sampling here focused on oxidation immediately post conventional ozonation and ozone nanobubble treatment, and after defined contact times to assess decay kinetics under high-load conditions. Figures 1h–1j show the lagoon-based treated wastewater system where samples were taken from the end of the process where treated wastewater was stored. This site was equipped with the first-of-a-kind portable ozone nanobubble generation units (Figure 1i) to enable flexible deployment and direct dosing into bloom-affected zones. Further detailed site information is provided in the supplementary information section (Table SI-1).

### 2.2. Ozonation and ozone nanobubble trials

Ozone nanobubble experiments were conducted using water samples collected directly from the ozone nanobubble system’s sampling port at each study site. Nanobubbles were generated by a Moleaer Lotus Nanobubble Generator which includes a 1.5HP self-priming pump with strainer basket, nanobubble generator and gas injection fitting, and a 20g/hr Moleaer ozone generator designed for the gas injection fitting. Samples were collected after ozone nanobubble injection into the water from a sampling port. These samples were retained for predetermined contact times (0, 5, 30, 60 and 90 min) to assess the progressive effects of treatment. In parallel, non-ozonated water was collected from the same water bodies, at the exact locations where feed water was drawn for the ozone nanobubble system, ensuring comparable source conditions. These samples were subjected to conventional ozonation locally, with similar ozone dosage (±5%) using a prepared ozone stock solution (supplementary information Table SI-2), enabling a direct comparison of treatment performance between ozone nanobubble application and conventional ozone exposure.

An ozone decay curve was monitored in a true-batch reactor for all samples (Table SI-2). The applied ozone doses were administered by injecting an aliquot of the ozone stock solution via a syringe into a continuously stirred 2-L glass reactor containing the sample and equipped with a floating Teflon lid to prevent ozone degassing. Ozone residuals (Zamyadi et al., 2015) were measured over a 90 min period by collecting 5 mL samples that were dispensed into 20 mL of indigo (0.02, 1 or 3%) solution; similar process was followed for ozone nanobubble samples. In parallel experiments, samples were collected for turbidity, total organic carbon (TOC) and dissolved organic carbon (DOC) measurements, cell viability, cell counts, total and dissolved toxin, and genomics analyses at predetermined contact times (0, 5, 30, 60 and 90 min). The samples (sampling volume of 150 mL) were immediately quenched with sodium bisulfite (NaHSO_3_ e 2.2 mg/mg O_3_) to stop the ozone reaction. The subsamples for dissolved toxins were immediately filtered over 0.45 µm filters and the filter permeates were stored in -25 °C for further analyses. The subsamples for total toxins were stored at -25 °C and were exposed to three phases of freeze-thaw to release all the cell-bound toxins prior to analyses. Bromate, the most commonly known ozonation by-product, was measured in all ozonated samples using 3,3’-dimethylnaphthidine as a sensing dye in photometry (Sigma-Aldrich Inc.). All samplings and analysis were conducted in triplicates, using the results to calculate the statistical significance and bias range (error bars in all graphs).

A modified indigo method was applied to quantify cumulative ozone diffusion from nanobubbles over time (Kuhn et al., 2025). This trial was conducted on three parallel (one with ozone nanobubble, one with conventional ozone, and one as control blank) 20L of water samples inoculated by cultured cells from Class-C sampling site. Immediately following ozonation nanobubble treatment, 90 mL of ozonated water was transferred to a 100 mL volumetric flask containing 10 mL of indigo reagent buffered to pH 2.5 with phosphoric acid. Samples were stored at 4 °C in the dark, and absorbance at 600 nm was measured every 30 minutes up to 90 minutes (maximum predetermined contact times for all the experiments). Parallel non-ozonated blanks were analysed to correct for background indigo degradation and matrix effects. Ozone concentration was calculated from the change in absorbance. This modified procedure captures both immediate and delayed ozone transfer from nanobubbles to the aqueous phase while minimizing interferences from peroxides and organic matter (Kuhn et al., 2025).

### 2.3. Cell counts, microscopic speciation, cell viability, cellular metabolite and water quality analysis

Algal and cyanobacterial species identification and enumeration on samples (100 mL sample volume) preserved in Lugol’s Iodine were conducted using a compound microscope and a Sedgewick Rafter counting chamber (Cooperative Research Centre for Freshwater Ecology, 1999). Colonies of *Microcystis* were also counted as they form colonies in real-world setting; in contrast to laboratory cultures which are mainly in separate cells. Water samples (100 mL sample volume) were analysed for both extracellular and total microcystins, cylindrospermopsin and anatoxins using ELISA kits (Chorus and Welker, 2021; Kibuye et al., 2021a). TOC and DOC measurements were conducted on an 820 Total Organic Carbon Analyser (Sievers Instruments Inc.). Measurements of UV absorbance at 254 nm (UV 254) were carried out on a UV1201 UV/vis Spectrophotometer (Shimadzu Corp.). Triplicate samples were filtered through 0.45 µm membrane filters prior to DOC and UV analyses.

The integrity of cyanobacterial cells in the full-scale treatment plants was evaluated (50 mL sample volume) using a fluorescein diacetate (FDA) and propidium iodide (PI) staining method for microscopy (Zamyadi et al., 2010; Zamyadi et al., 2019). The FDA stock solution (10 g/L) was prepared by dissolving FDA powder (Sigma-Aldrich) in a reagent grade acetone which was stored at -20 C until use. The FDA working solution (250 mg/L) was prepared by adding 25 mL of FDA stock solution into 1 mL of acetone. The FDA working solution was kept under refrigeration and used within 2 h after preparation to minimise degradation (Xi et al., 2011). The PI working solution (50 mg/L) was prepared by diluting 50 mL of PI solution (1 mg/mL, Sigma-Aldrich) in 1 mL of Milli-Q water and stored at 4 C until use (Xi et al., 2011). A simultaneous staining of the samples (Gumbo et al. 2014) was performed by adding an aliquot of 20 mL of FDA working solution (final concentration of 5.0 mg/L) and 133 mL of PI working solution to 1 mL of sub-sample in a 5 mL falcon tube (final concentration of 6.7 mg/L), and incubated at room temperature for 15 min, protected from light.

### 2.4. Genomic analysis

Samples collected for genomics analysis were filtered through 0.45 µm filter to concentrate the microbial contents of the water. Following filtration, each filter membrane was placed in 10 mL of the original water sample and vortexed for 2 mins to dislodge all microbes from the membrane. Sample concentrate was stored overnight at 4^°^C. For metagenomic sequencing 1 mL of the concentrate was extracted using a modification of the EZ2 PowerFecal Pro DNA/RNA Kit (QIAGEN). Sequencing libraries were prepared with Illumina DNA prep using quarter reagent reaction volumes and sequenced on the Illumina NextSeq2000 300 cycle kit (150bp PE). Schematic of metagenomic analysis pipeline is provided in supplementary information Table SI-3.

#### 2.4.1. Quality control, taxonomic profiling, metagenome assembly and ARG detection

Raw metagenomic reads were quality-filtered and trimmed using Fastp (v0.23.2 - Chen et al., 2018) with a minimum retained read length of 50 bp. Taxonomic classification of reads was performed using Kraken2 (v2.1.2 - Wood et al., 2019) with a confidence threshold of 0.1 against a database based on Genome Taxonomy Database (GTDB) release r214. Refined species-level abundance estimates were obtained using Bracken (Lu et al., 2017), and results were aggregated to phylum, family, and species levels. Only taxa with a mean relative abundance greater than 0.1% across samples were retained for downstream analyses.

Reads were assembled de novo using MEGAHIT (v1.1.3 - Li et al., 2015) and only contigs > 1000bp were retained. Contigs were taxonomically classified using Kraken2 as described above. Antimicrobial resistance genes were identified from assembled contigs using abricate (v1.0.1 - Seemann and Grüning, 2020) with the NCBI AMR database.

#### 2.4.2. Absolute abundance estimation

Quantitative metagenomic analysis was performed on T4 samples using the ZymoBIOMICS Spike-in Control II (Low Microbial Load) as an internal standard (Further details are provided in supplementary information Table SI-4). Bracken-estimated read counts for the three spike-in species were combined with manufacturer reported genome sizes and genome copy numbers to estimate sample-specific scaling factors. A robust linear model was fitted for each sample, regressing expected genome copy numbers against reads per base pair, with the intercept constrained to zero. This sample-specific scaling factor (units: genome copies per read per bp) was then applied to transform all microbial read counts into estimated absolute abundances. Genome sizes for non-spike-in species were based on median values for each species in GTDB. Data analysis was performed in R (v4.5.0) using the tidyverse packagen (v2.0.0) for data manipulation and plotting and MASS (v.7.3-65) for linear modelling.

## 3. Results and Discussion

Ozonation using nanobubble enhanced oxidation performance by 19%-34% in the drinking water reservoir compared to conventional ozonation while keeping the ozone dosage below 2mg/l. Figure 2a presents the comparative performance of conventional ozonation and ozone nanobubble treatment in reducing cyanobacterial biomass and cell viability in the drinking water reservoir across (locations shown in Figures 1a, 1b, 1c) seven sampling dates (beginning to end of the bloom season: 18 November, 6 January, 9 January, 23 January, 4 February, 14 February, and 7 March). The dominant cyanobacterial species during the study period was *Dolichospermum circinale*. Figure 2a shows the percentage reduction in total Cyanophyceae cell numbers at four time points post-dosing (5, 30, 60, and 90 minutes). While, aqueous ozone residual was below detection past 30 minutes of contact; however, modified indigo experiment have shown an additional 0.4 mg/L of accumulative ozone transfer in samples injected with nanobubble compared to conventional ozonation. These results are in accordance with Kuhn et al. (2025), demonstrating extended transfer of gaseous O_3_ from nanobubbles to the aqueous phase.

**Figure 2.**
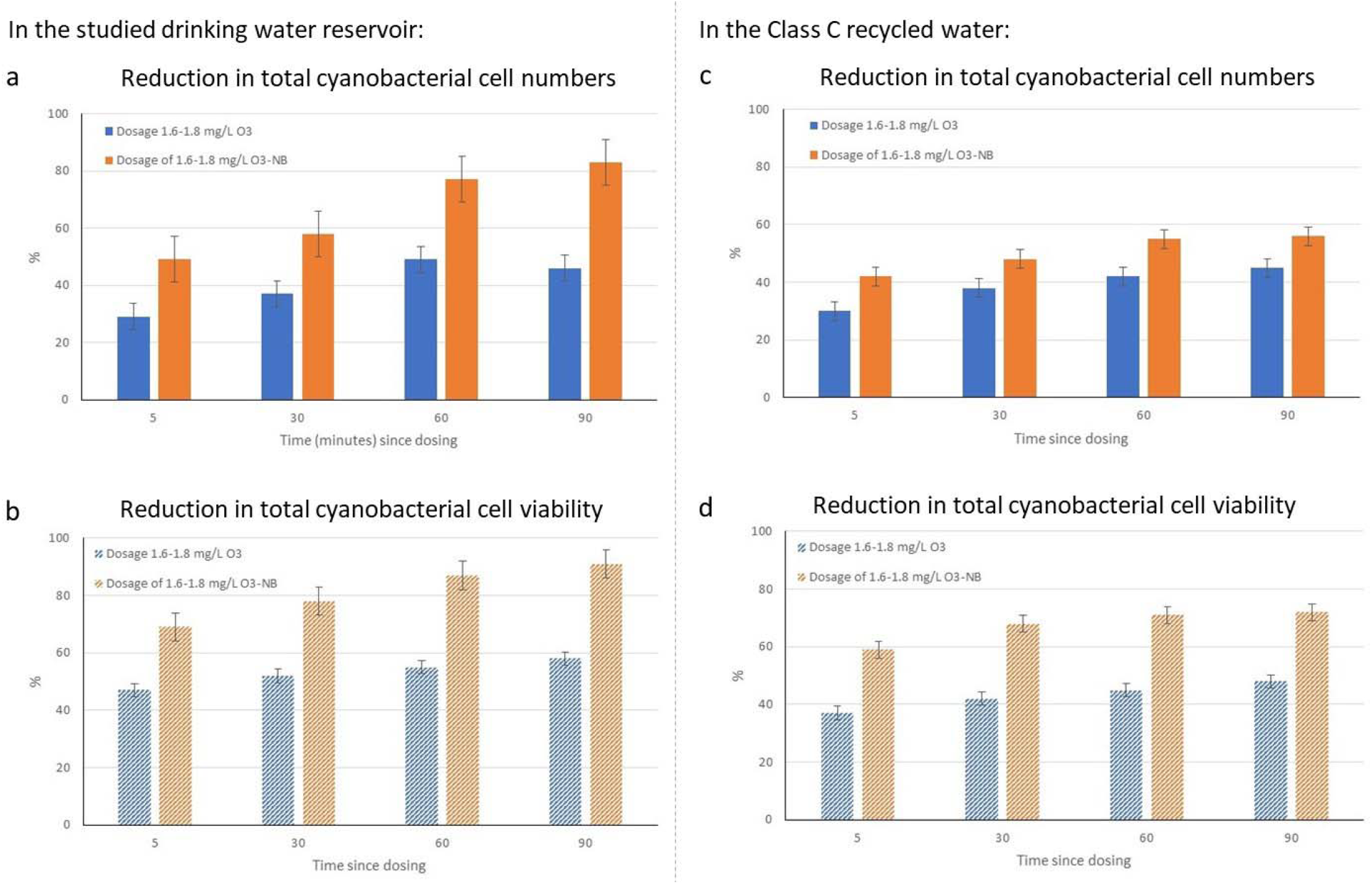
Comparative reduction of (a) cyanobacteria cell numbers and (b) total cell viability by conventional ozone and ozone nanobubbles in the studied water reservoir (error bars present variability between seven testing dates indicating repeatability of treatment performance.), and reduction of (c) eria cell numbers and (d) total cell viability by conventional ozone and ozone nanobubbles in the Class C recycled water (error bars present between four testing dates indicating repeatability of treatment performance).

During the entire experiment, ozone nanobubble consistently achieved higher removal rates than conventional ozone at the same initial dosing concentration (1.6–1.8 mg/L). For example, at 90 minutes, ozone nanobubble treatments achieved close to 90% reduction, compared to ∼50% for conventional ozone. The reproducibility of these results across multiple dates indicates stable and repeatable treatment performance under freshwater conditions, which is low organic and particulate loads (below 2 mg/L). In addition to cyanobacteria, the phytoplankton community composition included Bacillariophyceae, Chlorophyceae, Chrysophyceae, Cryptophyceae, Dinophyceae, and Euglenophyceae. Parallel analyses of these groups showed similar patterns, with ozone nanobubble delivering superior performance in reducing total cell counts across all classes.

The average cell lysis rates observed in this study for ozone nanobubble and conventional ozonation were, 0.97 ± 0.08 × 10^5^ M □ ^1^ s □ ^1^ and 0.86 ± 0.07 × 10^2^ M □ ^1^ s □ ^1^ accordingly. The *k*_*lysis*_ values for conventional ozonation are similar to a previous study on natural bloom samples by Zamyadi et al. (2015). Notably, the ozone nanobubble rates are three orders of magnitude larger than conventional ozonation; and despite being measured in natural water samples they are comparable to previous studies using cultured cyanobacteria (Wert et al., 2013; Ramseier et al., 2011). Cultured cyanobacteria samples provide more favourable oxidation conditions, with lower organic matter and a less diverse phytoplankton community background. Despite, the challenging nature of real-world conditions, the significantly enhanced performance observed with ozone nanobubbles is most likely due to their increased interfacial contact surface area, which enhance ozone–cell interactions beyond those achievable with conventional systems (Jia et al., 2023; Singh et al., 2024; Soyluoglu et al. 2021, 2022, 2023 and 2025; Chaffin et al., 2024; Koundle et al., 2024).

Figure 2b illustrates the reduction in total cell viability over the same time intervals and residual concentrations. Again, ozone nanobubble markedly outperformed conventional ozone, achieving ∼85–90% reduction in viable cells by 90 minutes, compared to ∼55–60% for conventional ozone. The more pronounced effect on cell viability than total cell counts suggests that ozone nanobubbles not only causes cell lysis but also effectively damaged any surviving cells, reducing the potential for regrowth. Measured toxins and bromate were below detection limit in all these samples.

Figure 2c compares the performance of conventional ozonation and ozone nanobubble treatment in reducing cyanobacterial biomass and viability in Class-C treated wastewater intended for recycling. Figure 2c shows the percentage reduction in total cyanobacterial cell numbers following four ozone and ozone nanobubble applications (T1: 11 November, T2: 17 December, T3: 6 January, and T4: 4 February) at a dosage of 1.6–1.8 mg/L, measured over four post-dosing intervals (5, 30, 60, and 90 minutes). At the beginning of the bloom season (T1: 11 November) *Pseudomonas* is dominant. By the height of the bloom season and summer heat (T2: 17 December, T3: 6 January, and T4: 4 February) *Microcystis* is dominant, showing a seasonal shift in dominant species. Across all time points, ozone nanobubble treatment achieved consistently higher reductions than conventional ozone, with improvements ranging from approximately 10–20% depending on the time interval. However, overall performance was lower compared to results from the drinking water reservoir trials (Figure 2a) due to the more complex wastewater matrix (significantly higher DOC than 4 mg/L), which likely increased ozone demand and reduced oxidative efficiency. This was reflected in lower oxidation rates as well (one order of magnitude lower than drinking water cases).

Figure 2d illustrates the reduction in total cell viability under the same treatment and dosage conditions. Again, ozone nanobubble markedly outperformed conventional ozone, achieving 60–75% reduction in viable cells compared to 35–45% for ozone alone. This greater reduction in viability relative to total cell numbers suggests that ozone nanobubbles effectively damage cell integrity and metabolic activity, even when complete lysis is not achieved. 64-71% reduction in total Microcystins were observed in these samples post ozonation with only 5% improvement by using ozone nanobubbles, this associated to oxidant reaction with high density cell biomass and associated organic matter (Lim et al., 2022). Bromate measurements were below detection limit in all these samples.

Genomics analysis provide the opportunity to study the performance of ozonation and the entire aquatic community reaction to treatment in more details compared to microscopic analysis (Zamyadi and Blackall, 2024). Figure 3a presents the relative abundance of microbial phylum across four time points (all measured at 90 minutes since dosage) from November to February, representing seasonal dynamics in treated wastewater samples subjected to ozone nanobubble oxidation. Each time point includes samples from two treatment stages, WWTP-post-aeration and Class C recycled water, both untreated and treated with ozonation. At the beginning of the bloom season (T1: 11 November), the microbial community was dominated by Pseudomonadota in untreated WWTP-post-aeration; while cyanobacteria were relatively abundant in Class C water as well. The application of ozonation at this stage resulted in a marked shift in Actinomycetota and Bacteroidota, with a concurrent increase in relative abundance of cyanobacteria and Bacteroidota, suggesting that ozone nanobubble treatment selectively suppressed some dominant groups.

**Figure 3.**
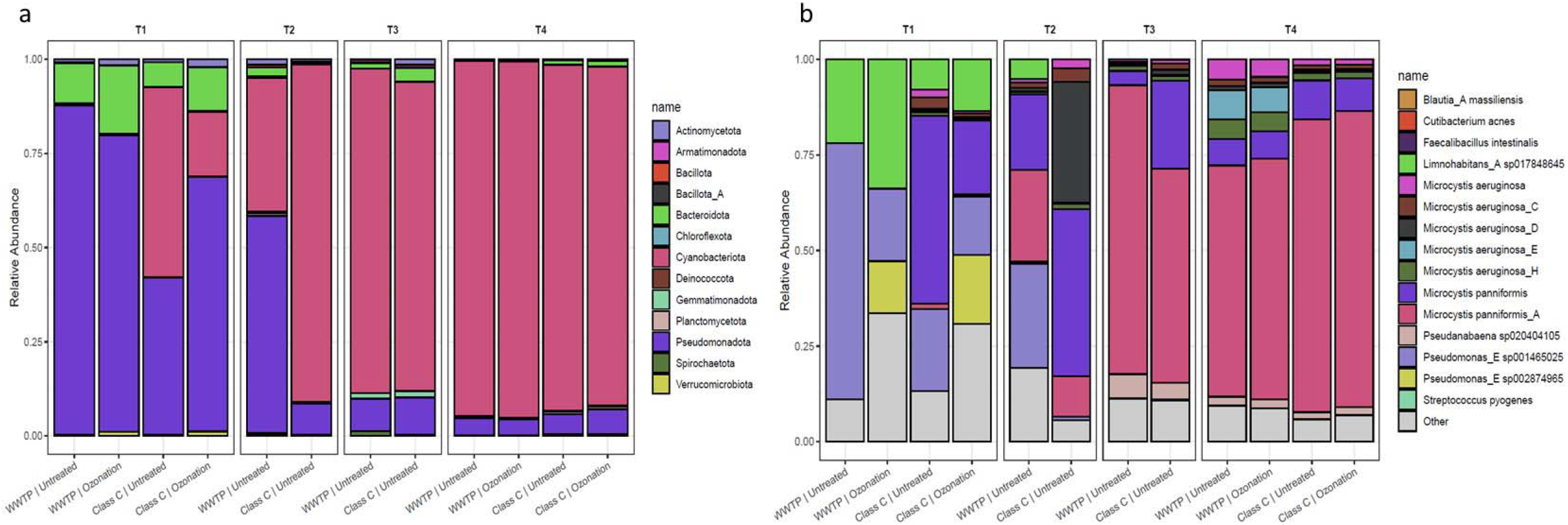
Dynamic composition of (a) total microbial and (b) cyanobacterial community with and without ozone nanobubble (90 minutes of contact) treatment in WWTP-post-aeration and Class-C recycled water (T1: 11 November, T2: 17 December, T3: 6 January, and T4: 4 February).

By December as the cyanobacterial bloom establishes with the hot weather, this shift became more pronounced, particularly in Class C recycled water, where ozonation led to a near-complete dominance of cyanobacteria in the remaining microbiome post ozonation. It is important to note that these results are presented as relative abundances, which reflect proportional changes in the community. To fully understand treatment effects, absolute abundances must also be considered in order to quantify the extent of cell lysis. This indicates a potential resilience of cyanobacterial blooms under certain oxidative conditions or seasonal nutrient dynamics. In contrast, un-ozonated WWTP-post-aeration and Class C recycled water retained a more mixed community, with Pseudomonadota, Bacteroidota, and other minor phyla still detectable. By mid-late summer (i.e. T3 and T4), the overall community composition across all samples, regardless of treatment, showed a clear dominance of cyanobacteria, exceeding 90% relative abundance, suggesting seasonal bloom pressure and/or oxidation tolerance in the dominant strains; which requires further investigation.

Despite this persistence, ozone nanobubble treatment consistently reduced the presence of several opportunistic or potentially resistant taxa, such as Bacteroidota, Pseudomonadota, and Gemmatimonadota, across all time points. The results suggest that while ozone nanobubble oxidation may not fully eliminate cyanobacterial taxa during peak bloom periods, it significantly restructures the microbial community by suppressing a broader range of bacterial contaminants. This reinforces the need to further investigate the role of ozone nanobubbles as a pre-treatment or polishing step in engineered water systems, particularly for seasonal bloom mitigation and community-level microbial control.

Figure 3b presents the relative abundance of cyanobacterial species across the same four seasonal time points (T1 to T4) in samples collected from WWTP-post-aeration and Class C recycled water, both untreated and following ozone nanobubble treatment. At the start of the bloom (i.e. T1), a diverse cyanobacterial community was observed, including the notable presence of *Pseudomonas* spp., *Microcystis* spp., and *Limnohabitans* spp. Ozone nanobubble oxidation at this early time point led to a visible shift in community composition, increasing dominance of select *Microcystis* and *Pseudanabaena* species, especially in Class C treated wastewater.

By December (i.e. T2), there was a pronounced increase in *Microcystis aeruginosa* and closely-related species (e.g., *M. aeruginosa*_*C, D*, and *E*), particularly in un-ozonated Class C water, indicating potential bloom onset and summer seasonal favourability for cyanobacterial proliferation. Ozone nanobubble treatment during this time appeared to reduce the diversity of *Pseudomonas* strains enhancing the growing dominance of *Microcystis* (in relative abundance). This can be due to strain-specific resistance, floc formation or environmental drivers which requires further investigation (Zamyadi and Blackall, 2024). Notably, *Microcystis panniformis* and its related variants remained relatively low in abundance but persisted across during all treatments.

By mid to late summer, the community shifted significantly toward dominance by *Microcystis* strains across all samples, regardless of ozone treatment. This seasonal bloom condition suggests strong environmental drivers, such as temperature or nutrient availability. However, ozone-treated samples still showed slightly lower diversity and reduced abundance of secondary opportunistic or potentially pathogenic taxa like *Pseudomonas* spp. These results underline the complexity of cyanobacterial bloom dynamics, particularly in treated wastewater stabilisation ponds (Romanis et al., 2023; Romanis et al., 2024), and highlight the need for early intervention strategies to control dominant strains before they reach bloom thresholds.

Figure 4 displays the true abundance (in genome copies /L) of various cyanobacterial species and co-occurring bacteria in different water matrices, following a controlled spike-in of target organisms. Unlike relative abundance plots, this chart reflects absolute genome copy counts, allowing for a more quantitative comparison of microbial load across sample types and treatment conditions. In WWTP-post-aeration, total abundance exceeded 2.0 × 10 □ genome copies /L, with *Microcystis aeruginosa* and its associated strains (*M. aeruginosa*_E, H, and p*anniformis*) dominating the community. After ozone nanobubble treatment, a reduction in total abundance was observed, yet dominant cyanobacterial species remained present in large numbers, indicating partial but not complete lysis under the applied conditions. A similar trend was noted in Class C recycled water: while spiked Class C samples showed high total abundance dominated by *Microcystis* strains, ozone nanobubble treatment resulted in a significant reduction, bringing total abundance below 1.0 × 10 □ genome copies /L (over 50% reduction), particularly affecting secondary species and low-abundance taxa.

**Figure 4.**
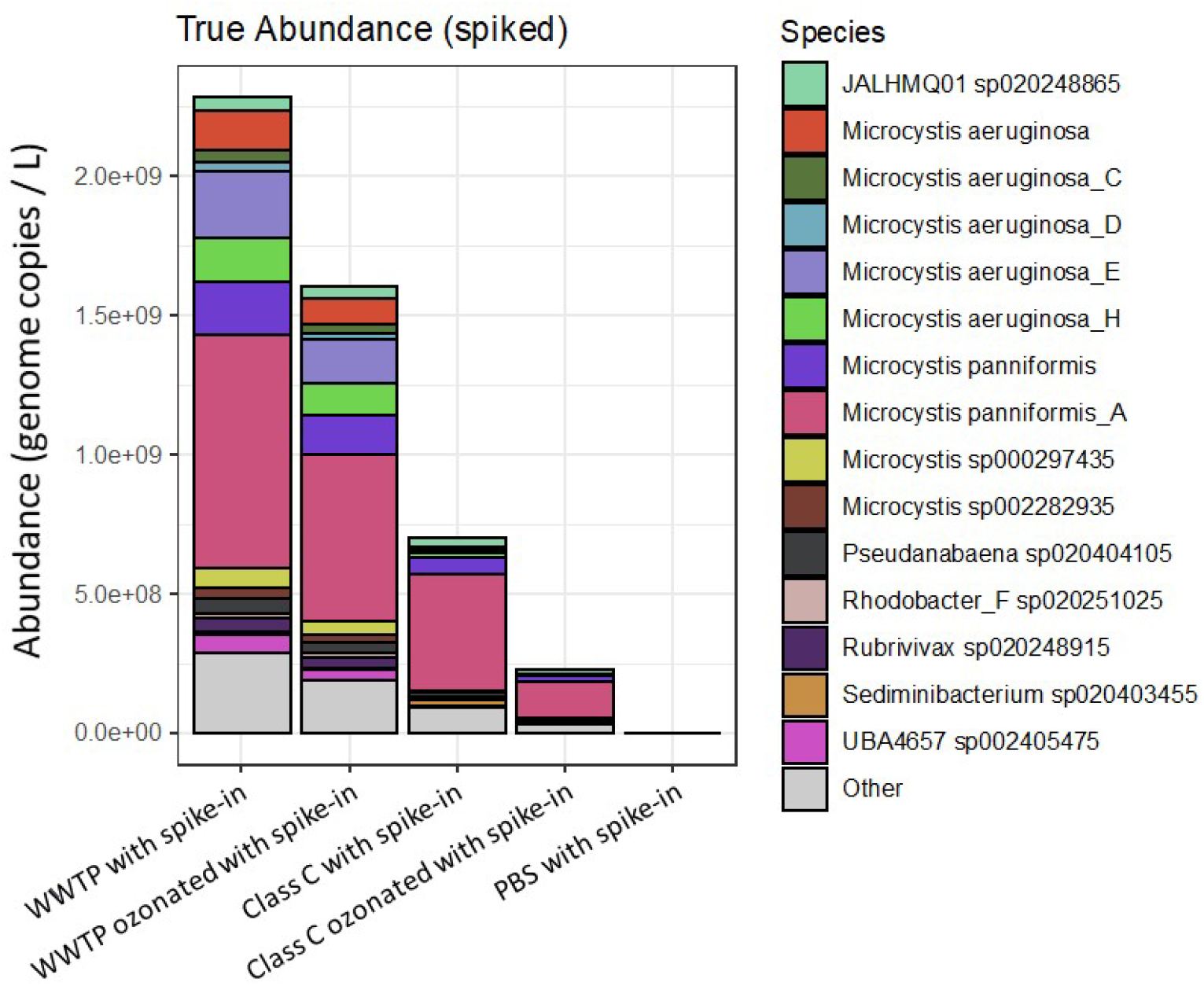
Absolute abundance of spiked cyanobacterial strains in WWTP-post-aeration and Class-C recycled water with and without ozone nanobubble oxidation (90 minutes of contact) at T4 (i.e. 4 February) only. “PBS with Spike in” is the extraction and sequencing negative control.

These genomics analyses provided the unique opportunity to investigate the fate of the full cyanobacterial bloom ecosystem, which includes the associated epibionts and antimicrobial resistance under ozone nanobubble oxidation and to explore knowledge gaps. Proposed mechanisms linking cyanobacterial blooms to ARG proliferation include the release of cyanobacterial metabolites that may facilitate horizontal gene transfer, the formation of phycospheres that create microenvironments favouring resistant bacterial communities, and the overall enrichment of ARGs as cyanobacterial biomass increases during bloom events (Zhang et al., 2020; Wang et al., 2020; Wang et al., 2021; Li et al., 2021; Wang et al., 2022; Volk et al., 2023; Ji et al., 2024). Additionally, cyanobacterial blooms are often characterized by dense and diverse microbial populations in close physical proximity, which may further enhance gene exchange. The eventual decline of blooms—potentially triggered by oxidation—could also lead to the sudden release of ARGs and other microbial contaminants, reinforcing the need for proactive and holistic treatment strategies at the source.

Detection of ARGs across the trial period was relatively limited, with only a few samples yielding positive results (Table 1). At the start of the bloom, ARGs were identified in both WWTP-post-aeration and Class C samples. These included *ere(D)*, an erythromycin resistance gene, and *blaPAU-1*, a beta-lactamase gene conferring resistance to β*-lactam* antibiotics. Interestingly, *ere(D)* was present both pre- and post-ozone nanobubble treatment in WWTP-post-aeration samples, suggesting that while oxidation may reduce viable bacterial cells, residual DNA from ARG carriers can persist in the water matrix. This persistence is consistent with studies such as Zheng et al. (2023) and Li et al. (2024), which found that ozonation can damage cells and degrade some extracellular DNA, but complete elimination of ARG sequences is challenging, particularly at lower oxidant doses or with high organic load.

**Table 1.**
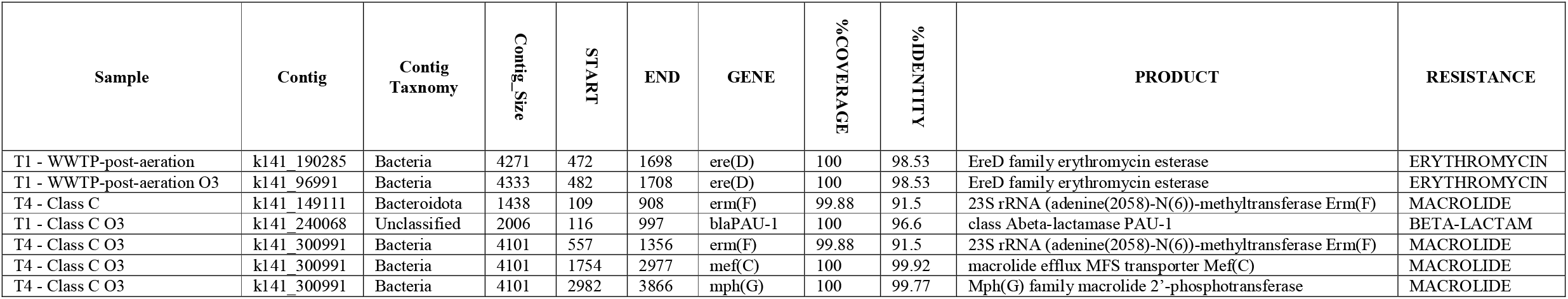
Results from ARG analysis pre and post ozone nanobubble application.

| Sample | Contig | Contig<br>Taxnomy | Contig_Size | START | END | GENE | %COVERAGE | %IDENTITY | PRODUCT | RESISTANCE |
| --- | --- | --- | --- | --- | --- | --- | --- | --- | --- | --- |
| T1 - WWTP-post-aeration | k141_190285 | Bacteria | 4271 | 472 | 1698 | ere(D) | 100 | 98.53 | EreD family erythromycin esterase | ERYTHROMYCIN |
| T1 - WWTP-post-aeration O3 | k141_96991 | Bacteria | 4333 | 482 | 1708 | ere(D) | 100 | 98.53 | EreD family erythromycin esterase | ERYTHROMYCIN |
| T4 - Class C | k141_149111 | Bacteroidota | 1438 | 109 | 908 | erm(F) | 99.88 | 91.5 | 23S rRNA (adenine(2058)-N(6))-methyltransferase Erm(F) | MACROLIDE |
| T1 - Class C O3 | k141_240068 | Unclassified | 2006 | 116 | 997 | blaPAU-1 | 100 | 96.6 | class Abeta-lactamase PAU-1 | BETA-LACTAM |
| T4 - Class C O3 | k141_300991 | Bacteria | 4101 | 557 | 1356 | erm(F) | 99.88 | 91.5 | 23S rRNA (adenine(2058)-N(6))-methyltransferase Erm(F) | MACROLIDE |
| T4 - Class C O3 | k141_300991 | Bacteria | 4101 | 1754 | 2977 | mef(C) | 100 | 99.92 | macrolide efflux MFS transporter Mef(C) | MACROLIDE |
| T4 - Class C O3 | k141_300991 | Bacteria | 4101 | 2982 | 3866 | mph(G) | 100 | 99.77 | Mph(G) family macrolide 2'-phosphotransferase | MACROLIDE |

By the establishment of the bloom (i.e. T4), ARG detection shifted towards macrolide resistance genes, particularly *erm(F), met(C)*, and *mph(G)*, identified in Class C water both pre- and post-ozonation (Table 1). These genes encode for methyltransferases and efflux transporters that protect bacteria from macrolide antibiotics. The recurrence of these genes post-ozone nanobubble treatment, despite overall microbial load reduction, suggests that oxidation did not fully eliminate either the resistant bacteria or extracellular DNA carrying these determinants. A persistent paradox in the literature is the apparent effectiveness of ARG inactivation in clean laboratory systems or with extracted DNA, contrasted with the relatively poor and inconsistent performance observed in complex, real-world water matrices (Lim et al., 2022). This discrepancy likely arises from multiple factors, including the physical location of ARGs (chromosomal vs. plasmid; intracellular vs. extracellular) and the lower-than-expected reactivity of DNA in situ. Furthermore, this observation aligns with recent work (Zheng et al., 2023; Li et al., 2024) showing that while advanced oxidation processes, including ozone nanobubbles, are effective at inactivating bacteria, complete removal of ARGs often requires higher oxidative stress or combined treatment approaches such as ozonation followed by biofiltration.

Figure 5 shows the performance of conventional ozonation and ozone nanobubble treatment in reducing *Microcystis*-dominated cyanobacterial biomass and viability in lagoon-based wastewater treatment systems, where initial *Microcystis* concentrations were very high. Across all time points (5, 30, 60, and 90 minutes post-dosing - Figure 5a), ozone nanobubble achieved greater reductions than conventional ozone—typically by 10–20%—but overall removal efficiencies were notably lower compared to treated wastewater or drinking water reservoir trials. At 90 minutes, ozone nanobubbles reduced cell counts by approximately 45– 50%, whereas conventional ozone achieved around 25–30%. This lower performance is attributed to the high particulate and organic load typical of lagoon-based wastewater treatment, which increases ozone demand and reduces the oxidant available for direct algal cell attack. 53-65% reduction in total Microcystins were observed in these samples post ozonation with 11% improvement by using ozone nanobubbles. Bromate measurements were below detection limit in all these samples.

**Figure 5.**
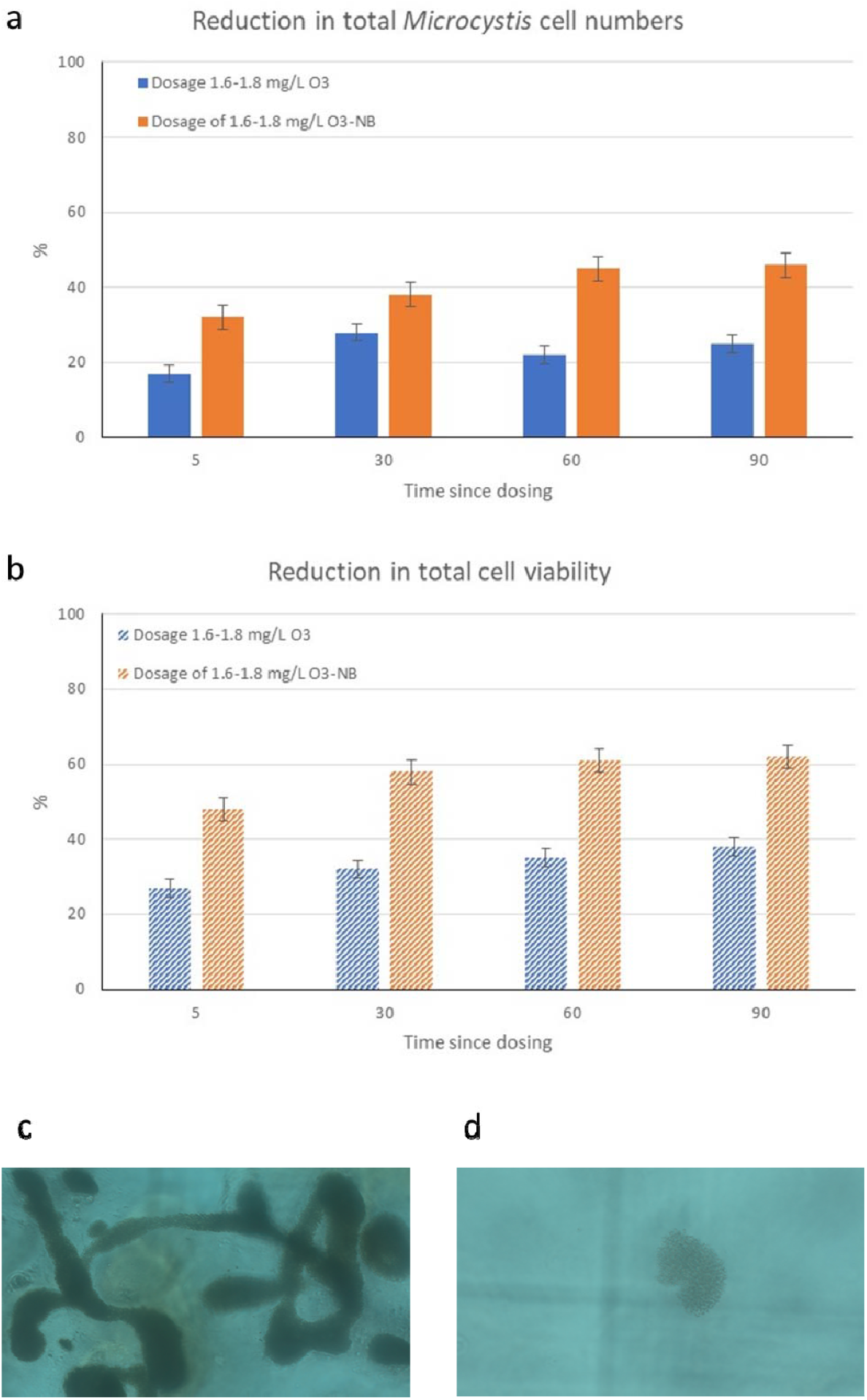
Comparative reduction of (a) cyanobacteria cell numbers and (b) total cell viability by conventional ozone and ozone nanobubbles in the lagoon-based wastewater treatment. Error bars present variability between two testing dates indicating repeatability of treatment performance. Figure c shows microscopic image of the water sample prior to ozone nanobubble application and post oxidation.

Figure 5b shows the reduction in *Microcystis* cell viability under the same conditions, where ozone nanobubble again outperformed conventional ozone, achieving 55–65% reduction compared to 25–40% for ozone alone. While the gains in viability reduction over conventional ozone were consistent, the absolute values remained modest due to the extreme cyanobacterial biomass and complex water matrix. The microscopic images (Figures 5c and 5d) illustrate typical *Microcystis* morphologies in these samples, including large colonies embedded in mucilaginous matrices, which likely offered significant protection against oxidant. Overall, these results highlight that although ozone nanobubble remains more effective than conventional ozone under challenging lagoon wastewater conditions, both treatments face limitations when *Microcystis* concentrations are extremely high in massive colonies with high self-shading and extracellular polymeric substance (EPS) protection.

## 4. Conclusion and Future Research Directions

This study demonstrates the potential of ozone nanobubble technology for mitigating cyanobacterial blooms, toxins, and co-occurring microbial contaminants across a range of engineered water systems. However, our findings also reveal several knowledge gaps and technical challenges that must be addressed before large-scale implementation:

1. Quantifying nanobubble persistence and distribution in complex water matrices: While the stability of ozone nanobubble is well-established under controlled laboratory conditions, their persistence and spatial distribution under real-world operational conditions remain unclear. In our trials, performance varied markedly between drinking water reservoir and high-organic treated wastewater matrices, likely reflecting differences in ozone demand, bubble–particle interactions, and ROS depletion. The development of nanobubble monitoring techniques—capable of resolving bubble size distribution, concentration, and lifetime in complex matrices—is needed to optimise dosing and placement strategies.
2. Understanding species- and strain-specific resistance to ozone nanobubble oxidation: These results showed that certain cyanobacteria, particularly *Microcystis aeruginosa* and *Dolichospermum circinale*, persisted post-treatment, especially in bloom-level biomass conditions. This persistence suggests that morphological and physiological traits, such as mucilaginous sheaths and colony structure, may confer protection from oxidative attack. Strain-level genomics and biochemical profiling, combined with oxidative stress assays, should be used to elucidate the mechanisms of resistance, ultimately guiding targeted dosing regimes or pre-treatment steps to overcome these protective traits.
3. Optimising ozone nanobubble dosing for dual control of cells and ARGs: Although ozone nanobubble treatment substantially reduced cyanobacterial biomass, ARGs including *ere(D), erm(F)*, and *blaPAU-1* were still detected post-oxidation in some samples. This indicates that cell inactivation alone does not ensure the degradation of associated DNA. Future studies should explore dose–response relationships for simultaneous removal of both viable cells and ARGs, evaluate the benefits of ROS enhancement, and assess the effectiveness of integrating post-oxidation barriers such as UV irradiation or biofiltration.
4. Evaluating DBP formation pathways unique to ozone nanobubble treatment: While bromate formation during conventional ozonation is well-characterised, and all measurements in this study were below detection limit, still little is known about DBP formation under ozone nanobubble treatment. The nano-scale gas–liquid interface and unique collapse dynamics of nanobubbles may influence radical yields and reaction pathways, particularly in bromide- or nitrogen-rich matrices. Long-term pilot studies should be conducted to characterise bromate and emerging DBPs in various source waters, enabling the development of safe and optimised operational envelopes for ozone nanobubble application.

These comparative trials across freshwater reservoirs, treated wastewater for recycling, and lagoon-based systems revealed that ozone nanobubble efficacy is strongly matrix-dependent, with highest performance in low-organic waters. The ability to quantify community shifts at both relative and absolute levels, down to strain resolution, represents a significant advance in monitoring and process optimization, ensuring targeted, evidence-based improvements in water safety and treatment efficacy.

Together, these results reinforce the value of ozone nanobubble technology as a promising yet not standalone strategy for bloom control and microbial risk reduction in both drinking water and wastewater reuse systems. Compared to conventional ozonation, ozone nanobubble demonstrated extended oxidative activity, reduced cell viability, and better retention of performance at lower ozone doses. However, full control of cyanobacterial outbreaks— especially under peak bloom conditions—will likely require integration of ozone nanobubble with complementary strategies such as powdered activate carbon, biological filtration, or early detection-based dosing.

Scaling ozone nanobubble systems to full-scale treatment plants requires further investigation, including modelling energy and operational costs, and evaluating seasonal performance under variable contaminant loads.

## Acknowledgment

The authors acknowledge the support from the water industry partner, the Australian Research Council Discovery project #DP250101804, Water Research Australia project #4554, Mallee Regional Innovation Centre (MRIC) which is a partner in the Victoria Drought Resilience Adoption and Innovation Hub, Water Research Foundation project #5237, and Monash University Department of Civil and Environmental Engineering and Faculty of Engineering academic establishment funds.

## References

Atkinson, A.J., Apul, O.G., Schneider, O., Garcia-Segura, S., Westerhoff, P. (2019) Nanobubble technologies offer opportunities to improve water treatment. Accounts of Chemical Research, 52, 5, 1196–1205.

Batagoda, J.H., Hewage, S.D. A., Meegoda, J.N. (2019). Nano-ozone bubbles for drinking water treatment. Journal of Environmental Engineering and Science, 14(2), 57–66.

Chaffin, J.D., Berthold, D.E., Braig, E.C., Fuchs, J.D., Gabor, R.S., Jacquemin, S.J., Kuhn, H.E., Labus, L.D., Laughinghouse, H.D., Lefler, F.W., Mash, H.E., Raymond, H.A., Stanley, H., Taylor, A.T., Weavers, L.K., Wendel., S. (2024). Effectiveness of ozone nanobubble treatments on high biomass cyanobacterial blooms: A mesocosm experiment and field trial. Journal of Environmental Management, 372, 123406.

Chen, C., Zamyadi, A., Lin, T.-F., Gallagher, D. (2019). Removal of odourants from drinking water. In T.-F. Lin, S. Watson, & I. H. Suffet (Eds.), Taste and Odour in Source and Drinking Water: Causes, Controls, and Consequences. IWA Publishing.

Chen, S., Zhou, Y., Chen, Y., Gu, J. (2018) fastp: an ultra-fast all-in-one FASTQ preprocessor. Bioinformatics, 34 (17), i884–i890.

Chorus, I., Welker, M. (2021) Toxic Cyanobacteria in Water, 2nd edition. CRC Press, Boca Raton. (FL) USA, on behalf of the World Health Organization (WHO).

Coggins, L.X., Crosbie, N.D., Ghadouani, A. (2019) The small, the big, and the beautiful: Emerging challenges and opportunities for waste stabilization ponds in Australia. WIREs Water, 6.

Coral, L.A., Zamyadi, A., Berbeau, B., Bassetti, F.J., Lapolli, F.R., Prevost, M. (2013) Oxidation of Microcystis aeruginosa and Anabaena flos-aquae by ozone: Impacts on cell integrity and chlorination by-product formation. Water Research, 47(9), 2983–2994.

Gumbo, J.R., Cloete, T.E., van Zyl, G.J.J., Sommerville, J.E.M. (2014). The viability assessment of Microcystis aeruginosa cells after co-culturing with Bacillus mycoides B16 using flow cytometry. Phys. Chem. Earth 72e75, 24e33.

Ikehata, K., Li, Y. (2018) Ch 5: Ozone-Based Processes; In Advanced Oxidation Processes for Waste Water Treatment Emerging Green Chemical Technology, 2018, Pages 115–134.

Ji, W., Ma, J., Zheng, Z., Al-Herrawy, A.Z., Xie, B., Wu, D. (2024) Algae blooms with resistance in fresh water: Potential interplay between Microcystis and antibiotic resistance genes. Science of the Total Environment, 940 (2024) 173528.

Jia, M., Farid, M.U., Kharraz, J.A., Kumar, N.M., Chopra, S.S., Jang, A., Chew, J., Khanal, S.K., Chen, G., An, A.K. (2023) Nanobubbles in water and wastewater treatment systems: Small bubbles making big difference. Water Research, 245, 120613.

Kibuye, F.A., Almuhtaram, H., Zamyadi, A., Gaget, V., Owen, C., Hofmann, R., Wert, E.C. (2021a) Utility practices and perspectives on monitoring and source control of cyanobacterial blooms. AWWA Water Science (JAWWA), 3 (6), e1264.

Kibuye, F.A., Zamyadi, A., Wert, E.C. (2021b) A critical review on operation and performance of source water control strategies for cyanobacterial blooms: Part I-Chemical control methods. Harmful Algae. 109, 102099.

Kibuye, F.A., Zamyadi, A., Wert, E.C. (2021c) A critical review on operation and performance of source water control strategies for cyanobacterial blooms: Part II-Mechanical and biological control methods. Harmful Algae. 109, 102119.

Koundle P., Nirmalkar N., Momotko M., Boczkaj G. (2024). Ozone nanobubble technology as a novel AOPs for pollutants degradation under high salinity conditions. Water Research, 263, 122148.

Krienitz, L., Dadheech, P. K., Fastner, J., Kotut, K. (2013). The rise of potentially toxin producing cyanobacteria in Lake Naivasha, Great African Rift Valley, Kenya. Harmful Algae, 27, 42–51.

Kuhn, H.E., Fagan, W.P., Fuchs, J.D., Gabor, R.S., Raymond, H.A., Weavers, L.K. (2025 - In-press). Modified indigo method to quantify ozone diffusion from nanobubbles. Ozone: Science & Engineering.

Li, D., Liu, C-M., Luo, R., Sadakane, K., Lam, T-W. (2015) MEGAHIT: an ultra-fast singlenode solution for large and complex metagenomics assembly via succinct de Bruijn graph. Bioinformatics, 31(10), 1674–1676.

Li, W., Mao, F., Te, S.H., He, Y., Gin, K.Y.-H. (2021) Impacts of Microcystis on the dissemination of the antibiotic resistome in cyanobacterial blooms. ACS EST Water, 1, 1263−1273.

Li, F., Liu, K., Bao, Y., Li, Y., Zhao, Z., Wang, P., Zhan, S. (2024) Molecular level removal of antibiotic resistant bacteria and genes: A review of interfacial chemical in advanced oxidation processes. Water Research, 254, 121373.

Lim, S., Shi, J.L., von Gunton, U., McCurry, D.L. (2022) Ozonation of organic compounds in water and wastewater: A critical review. Water Research, 213, 118053

Lu, J., Breitwieser, F.P., Thielen, P., Salzberg, S.L. (2017) Bracken: estimating species abundance in metagenomics data. PeerJ Computer Science, 3:e104.

Manasfi, T. (2021). Chapter Four - Ozonation in drinking water treatment: an overview of general and practical aspects, mechanisms, kinetics, and byproduct formation. Comprehensive Analytical Chemistry. T. Manasfi and J.-L. Boudenne, Elsevier. 92: 85–116.

Merel, S., Walker, D., Chicana, R., Snyder, S., Baurès, E., & Thomas, O. (2013). State of knowledge and concerns on cyanobacterial blooms and cyanotoxins. Environment International, 59, 303–327.

Morrison, C., A. Atkinson, A. Zamyadi, F. Kibuye, M. McKie, S. Hogard, P Mollica, S. Jasim, Eric C. Wert (2021) Critical review and research needs of ozone applications related to virus inactivation: Potential implications for SARS-CoV-2. Ozone: Science & Engineering, 43(1), 2–20.

Paerl, H. W., & Paul, V. J. (2012) Climate change: Links to global expansion of harmful cyanobacteria. Water Research, 46(5), 1349–1363.

Ramseier, M.K., von Gunten, U., Freihofer, P., Hammes, F. (2011) Kinetics of membrane damage to high (HNA) and low (LNA) nucleic acid bacterial clusters in drinking water by ozone, chlorine, chlorine dioxide, monochloramine, ferrate(VI), and permanganate. Water Research, 45, 1490e1500.

Romanis, C.S., Timms, V.J., Nebauer, D.J., Crosbie, N.D., Neilan, B.A. (2023) Microbiome analysis reveals Microcystis blooms endogenously seeded from benthos within wastewater maturation ponds. Applied & Env Microb, 90.

Romanis, C.S., Timms, V.J., Crosbie, N.D., Neilan, B.A. (2024) 16S rRNA gene amplicon sequencing data from an Australian wastewater treatment plant. Microbiology Resource Announcements, 13.

Seemann, T., Grüning, B. (2020) https://github.com/tseemann/abricateIssue#138.

Sgroi, M., Roccaro, P., Oelker, G., Snyder, S.A. (2016) N-nitrosodimethylamine (NDMA) formation during ozonation of wastewater and water treatment polymers. Chemosphere, 144, 1618–1623.

Singh, E., Kumar, A., Lo, S.-L. (2024) Advancing nanobubble technology for carbon-neutral water treatment and enhanced environmental sustainability. Environmental Research, 252(3), 118980.

Soyluoglu, M., Kim, D., Zaker, Y., Karanfil, T. (2021) Stability of oxygen nanobubbles under freshwater conditions. Water Research, 206, 117749.

Soyluoglu, M., Kim, D., Zaker, Y., Karanfil, T. (2022) Removal mechanisms of geosmin and MIB by oxygen nanobubbles during water treatment. Chemical Engineering Journal, 443, 136535

Soyluoglu, M., Kim, D., Zaker, Y., Karanfil, T. (2023) Characteristics and stability of ozone nanobubbles in freshwater conditions. Environ. Sci. Technol., 57(51), 21898–21907st

Soyluoglu, M., Williams, C. F., Karanfil, T. (2025) Advancing water treatment: Ozone nanobubbles for geosmin and 2-methylisoborneol control. Chemical Engineering Journal, 513, 162892.

Su, M., Yu, J., Zhang, J., Chen, H., An, W., Vogt, R.D., Andersen, T., Jia, D., Wang, J, Yang, M. (2015) MIB-producing cyanobacteria (Planktothrix sp.) in a drinking water reservoir: Distribution and odor producing potential. Water Research, 68, 444–453.

Su, M., Suruzzaman, MD., Zhu, Y., Lu, J., Yu, J., Zhang, Y., Yang, M. (2021) Ecological niche and in-situ control of MIB producers in source water. Journal of Environmental Sciences, 110, 119–128.

Volk, A., Lee, J. (2023) Review article - Cyanobacterial blooms: A player in the freshwater environmental resistome with public health relevance? Environmental Research, 216, 114612.

Wang, B., Peng, Q., Wang, R., Yu, S., Li, Q., & Huang, C. (2022). Efficient Microcystis removal and sulfonamide-resistance gene propagation mitigation by constructed wetlands and functional genes analysis. Chemosphere, 292, 133481.

Wang, Z., Chen, Q., Zhang, J., Guan, T., Chen, Y., & Shi, W. (2020). Critical roles of cyanobacteria as reservoir and source for antibiotic resistance genes. Environment International, 144, 106034.

Wang, Z., Chen, Q., Zhang, J., Yan, H., Chen, Y., Chen, C., & Chen, X. (2021). High prevalence of unstable antibiotic heteroresistance in cyanobacteria causes resistance underestimation. Water Research, 202, 117430.

Wert, E.C., Dong, M.M., Rosario-Ortiz, F.L. (2013). Using digital flow cytometry to assess the degradation of three cyanobacteria species after oxidation processes. Water Research, 47, 3752e3761.

Wood, D.E., Lu, J., Langmead, B. (2019) Improved metagenomic analysis with Kraken 2. Genome Biology, 20, 257.

Xi, X., Zhi-ying, H., Ying-xu, C., Xin-qiang, L., Hua, L., Yi-chao, Q. (2011). Optimization of FDA-PI method using flow cytometry to measure metabolic activity of the cyanobacteria, Microcystis aeruginosa. Phys. Chem. Earth 36, 424e429.

Yang, X., Chen, L., Oshita, S., Fan, W., Liu, S. (2023). Mechanism for enhancing the ozonation process of micro-and nanobubbles: bubble behavior and interface reaction. ACS EST Water, 3(12), 3835–3847.

Zamyadi, A., Ho, L., Newcombe, G., Daly, R.I., Burch, M., Baker, P., Prevost, M. (2010). Release and oxidation of cell-bound saxitoxins during chlorination of Anabaena circinalis cells. Environ. Sci. Technol. 44, 9055e9061.

Zamyadi, A., MacLeod, S. L., Fan, Y., McQuaid, N., Dorner, S., Sauvé, S., & Prévost, M. (2012). Toxic cyanobacterial breakthrough and accumulation in a drinking water plant: A monitoring and treatment challenge. Water Research, 46(5), 1511–1523.

Zamyadi, A., L. A. Coral, B. Barbeau, S. Dorner, F. R. Lapolli, M. Prévost (2015) Fate of toxic cyanobacterial genera from natural bloom events during ozonation. Water Research, 73, 204–215.

Zamyadi A., Romanis C., Mills T., Neilan B., Choo F., Coral L., Gale D., Newcombe G., Crosbie N., Stuetz R., Henderson R. (2019) Diagnosing water treatment critical control points for cyanobacterial removal: Exploring benefits of combined microscopy, next-generation sequencing, and cell integrity methods. Water Research, 152, 96–105.

Zamyadi A., Glover C.M., Yasir A., Stuetz R., Newcombe G., Crosbie N.D., Lin T-F., Henderson R. (2021) Toxic cyanobacteria in water supply systems: data analysis to map global challenges and demonstrate the benefits of multi-barrier treatment approaches. H2Open Journal, 4 (1): 47–62.

Zamyadi A., Blackall L. (2024) Harmful and nuisance blue-green algae in water supply and sludge/washwater recovery systems: Investigating accumulation phenomenon and management strategies. Mallee Regional Innovation Centre (MRIC), Mildura, Victoria, Australia

Zhang, Q., Zhang, Z., Lu, T., Peijnenburg, W. J. G. M., Gillings, M., Yang, X., Chen, J., Penuelas, J., Zhu, Y.-G., Zhou, N.-Y., Su, J., & Qian, H. (2020). Cyanobacterial blooms contribute to the diversity of antibiotic-resistance genes in aquatic ecosystems. Communications Biology, 3(1), 1–10.

Zheng, Q., Zhang, Y., Zhang Q., Wang, Y., Yu, G. (2023) Removal of antibiotic resistant bacteria and plasmid-encoded antibiotic resistance genes in water by ozonation and electro-peroxone process. Chemosphere, 319, 138039.

Zhou, J., Qu, M., Dunkinson, C., Lefebvre, D.D., Wang, Y., Brown, R.S (2023) The Effect of Microcystis on the Monitoring of Faecal Indicator Bacteria. Toxins, 15, 628.

