## Supplementary information for "Exploring use of ozone nanobubbles for removal of cyanobacteria and co-occurring antimicrobial resistance genes in water supply and reuse systems"

Table SI-1. Detailed site information (maps sourced from Google Maps 2025)

| “Yellow star” shows location of experiment at the drinking water reservoir:  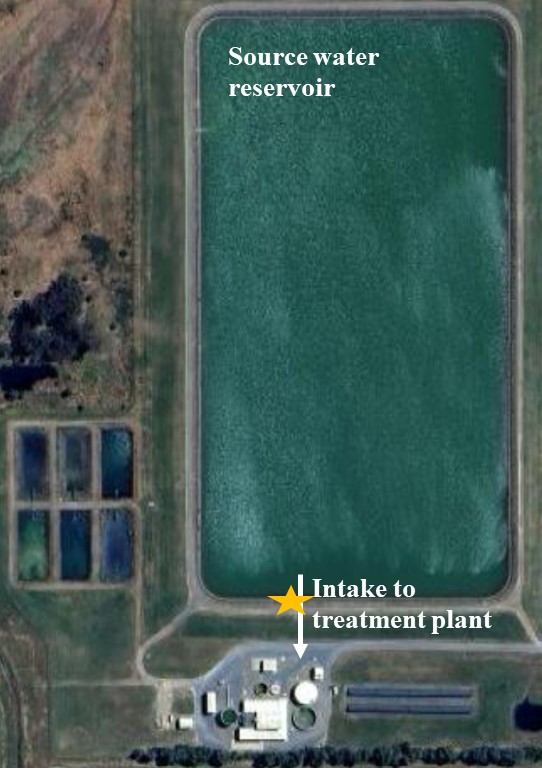 | “Yellow star 1” shows location of experiment for treated wastewater post aeration ponds, labelled “WWTP-aeration-pond”, and “Yellow start 2” show location of experiment for treated wastewater at the end of storage ponds before recycled for farming labelled “Class-C”:  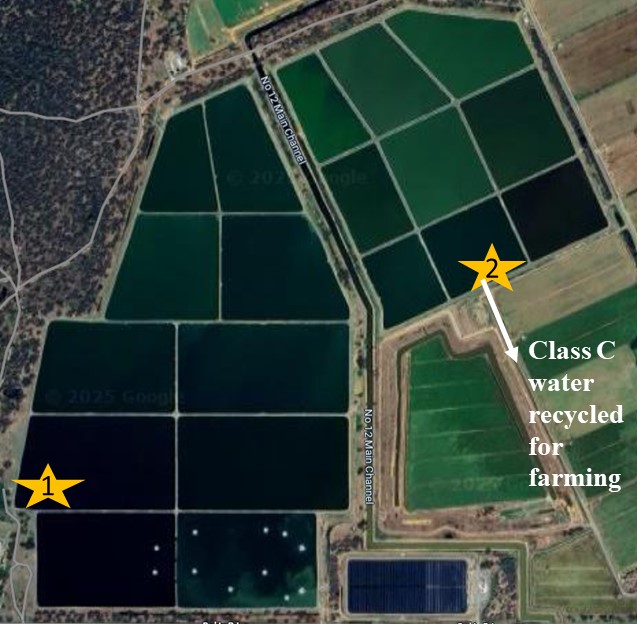 | Lagoon-based wastewater treatment plant “Yellow star” shows location of the experiment:  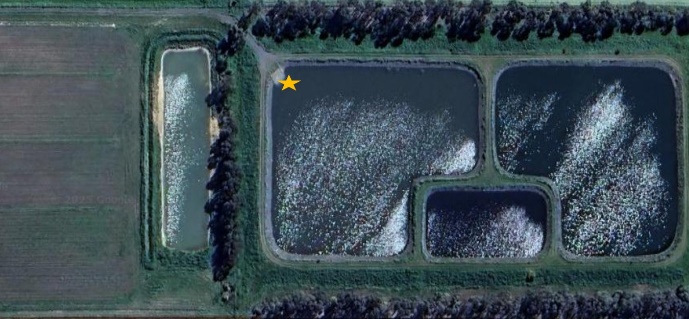 |
| --- | --- | --- |

Table SI-2. Ozone stock solution preparation and decay (Adapted from Zamyadi et al., 2015)

| Ozone stock solution | Stock ozone solutions (60–70 mg O₃ L⁻¹) were freshly prepared prior to each experimental run. Gaseous ozone was generated using a bench-top ozone generator (Model TG-10, Ozone Solutions Inc.) and continuously diffused into a 1 L borosilicate glass flask containing ultrapure water (Milli-Q, 18.2 MΩ cm) maintained at 4 °C to enhance ozone solubility and stability. The ozonated water was allowed to equilibrate until the target concentration was achieved, as determined by spectrophotometric measurement.  Ozone concentrations in both the stock solution and experimental samples were quantified following the Standard Methods for the Examination of Water and Wastewater, Method 4500-O₃ (APHA, 2005). The indigo trisulfonate colorimetric method was applied, in which the decrease in absorbance of the indigo dye at λ = 600 nm (molar extinction coefficient ε = 20,000 M⁻¹ cm⁻¹) is proportional to the dissolved ozone concentration. Absorbance was measured using a UV–Vis spectrophotometer (Cary 100, Varian) equipped with 1 cm and 2 cm quartz cuvettes, depending on sample concentration and absorbance range.  All ozone measurements were performed immediately after sample collection to minimise ozone decay. Dissolved ozone residuals were reported as mean values from triplicate measurements, with standard deviations less than 5% of the mean. |
| --- | --- |
| Ozone decay curve | Ozone is inherently unstable in aqueous environments, undergoing spontaneous decomposition influenced by temperature, pH, and the presence of organic and inorganic constituents. To characterise ozone persistence under experimental conditions, an ozone decay curve was established for each treatment scenario. Measurements were conducted in a true-batch reactor system corresponding to each combination of ozone dose and initial cell density. Ozone concentration was monitored at predetermined time intervals until complete decay was observed, enabling the determination of both ozone half-life and effective exposure duration for subsequent oxidation analyses. |

Table SI-3. Schematic of metagenomic analysis pipeline (Li et al., 2015; Lu et al., 2017; Chen et al., 2018; Wood et al., 2019; Seemann and Grüning, 2020; Maire et al., 2024)

| Metagenomic analysis pipeline | 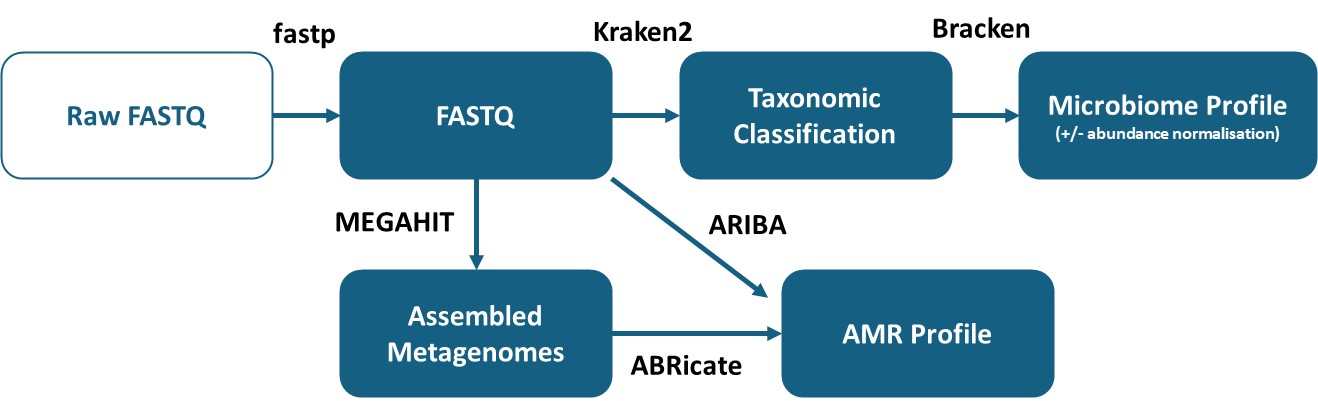 |
| --- | --- |

Table SI-4. Details of quantitative metagenomic analysis

| ZymoBIOMICS Spike-in Control II | 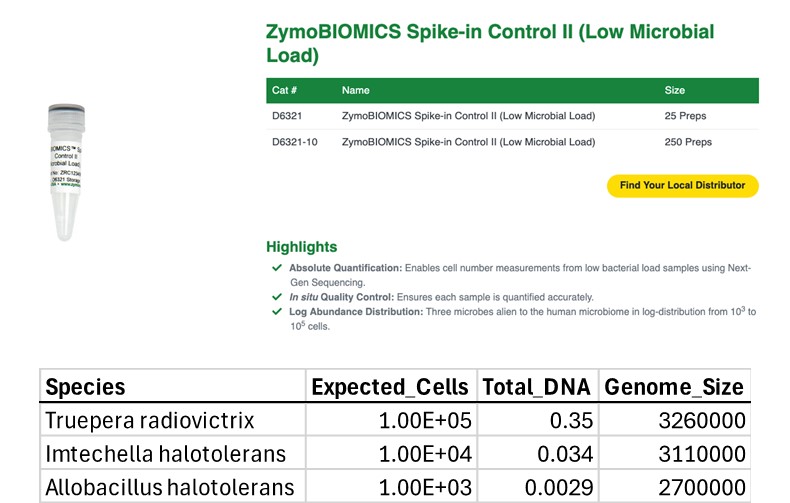 |
| --- | --- |
| Testing quantification | 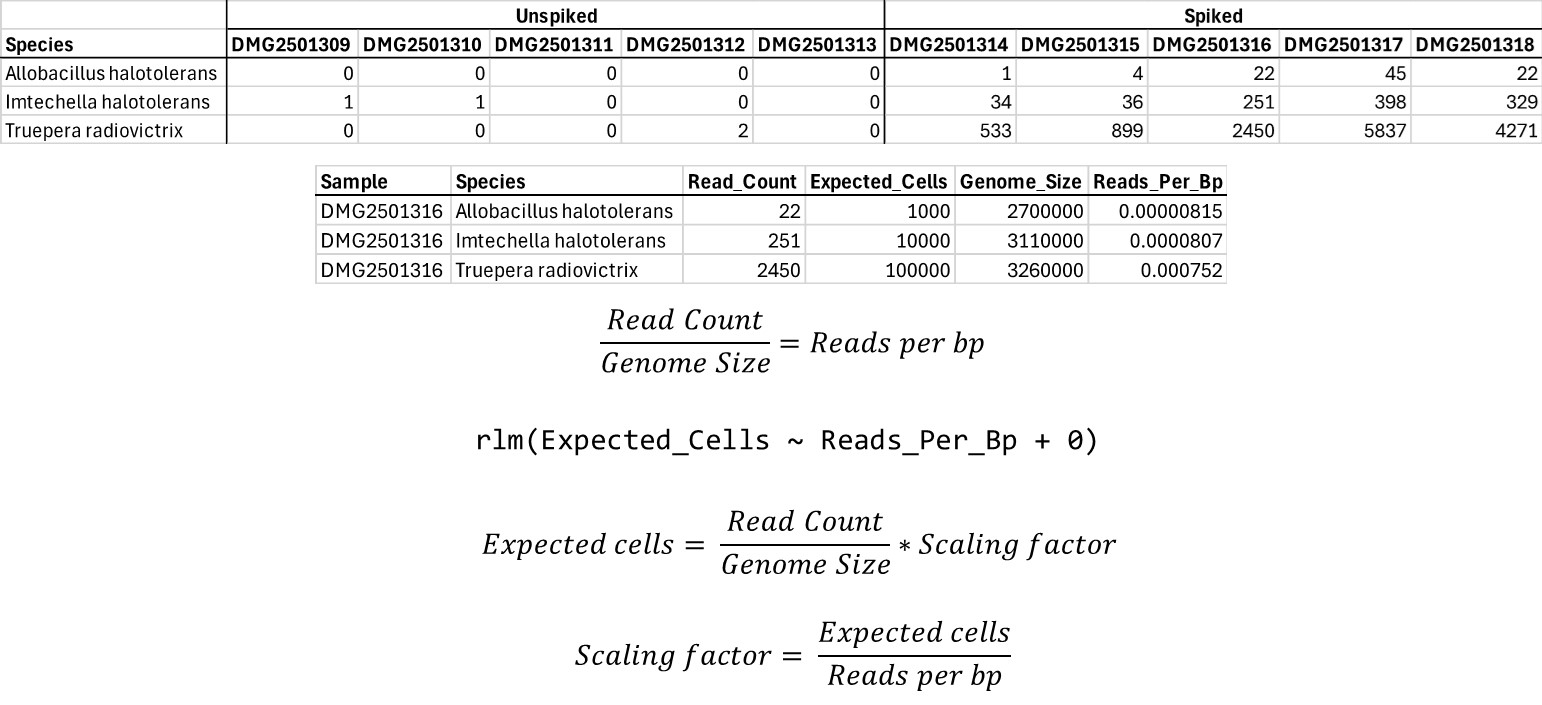 |

**References**

American Public Health Association (APHA), 2005. Standard Methods for the Examination of Water and Wastewater, twenty first ed. American Public Health Association (AWWA) and the Water Environment Federation, Washington, DC, USA.

Chen, S., Zhou, Y., Chen, Y., Gu, J. (2018) fastp: an ultra-fast all-in-one FASTQ preprocessor. *Bioinformatics*, 34 (17), i884–i890.

Li, D., Liu, C-M., Luo, R., Sadakane, K., Lam, T-W. (2015) MEGAHIT: an ultra-fast single-node solution for large and complex metagenomics assembly via succinct de Bruijn graph. *Bioinformatics*, 31(10), 1674–1676.

Lu, J., Breitwieser, F.P., Thielen, P., Salzberg, S.L. (2017) Bracken: estimating species abundance in metagenomics data. *PeerJ Computer Science*, 3:e104.

Seemann, T., Grüning, B. (2020) https://github.com/tseemann/abricate Issue#138.

Maire, J., Collingro, A., Tandon, K., Jameson, V.J., Judd, L.M., Horn, M., Blackall, L.L., van Oppen, M.J.H. (2024) Chlamydiae as symbionts of photosynthetic dinoflagellates. *The ISME Journal*, 18(1), wrae139.

Wood, D.E., Lu, J., Langmead, B. (2019) Improved metagenomic analysis with Kraken 2. *Genome Biology*, 20, 257.

Zamyadi, A., L. A. Coral, B. Barbeau, S. Dorner, F. R. Lapolli, M. Prévost (2015) Fate of toxic cyanobacterial genera from natural bloom events during ozonation. *Water Research*, 73, 204-215.
